# A stereotyped glial attachment determines the morphology and function of neuronal cilia

**DOI:** 10.64898/2026.04.08.717278

**Authors:** Leland R. Wexler, Brian Griffin, Priya Dutta, Piali Sengupta, Irina Kolotuev, Niels Ringstad, Maxwell G. Heiman

## Abstract

Primary cilia are signaling hubs that influence nearly every aspect of cell physiology. Neuronal primary cilia were recently discovered to interact with glia in the mammalian brain, but how cilia-glia attachments form and what roles they play remain unknown. Here, we find that two *C. elegans* sensory neurons (URX and BAG) use their primary cilia to attach to a specific glial partner (ILsoL). Through a genetic screen, we find that cilia-glia attachment requires BUG-1, a secreted protein with conserved cell adhesion domains that localizes to the cilia-glia attachment site. In the absence of BUG-1, neuronal cilia are present but fail to attach to the glial cell. We find that loss of cilia-glia attachment alters cilia morphology and disrupts stimulus-evoked calcium dynamics in cilia. We propose that primary cilia not only act as antennae for long-range cell communication, but also form close-range cell attachments that modulate cell signaling.

## INTRODUCTION

Primary cilia are found on neurons and glia throughout the brain, where they play critical but poorly understood roles in neural development and function. For example, genetic disorders affecting cilia (ciliopathies) cause severe neurological phenotypes whose etiology is unknown^1^. Alterations in ciliary gene expression are associated with neuropsychiatric and neurodegenerative disorders, including autism, schizophrenia, bipolar disorder, Parkinson’s disease, and amyotrophic lateral sclerosis, but the underlying causal mechanisms have not been determined^2–7^. Understanding how primary cilia contribute to neuronal health and disease will require determining the neuron-specific functions of cilia and defining the molecular mechanisms that support those functions.

Recent ultrastructural studies of neuronal primary cilia in the mammalian brain revealed that cilia form intimate contacts with both neighboring neurons and glia within a dense extracellular environment^8–10^. Some cilia are apposed to release sites for synaptic vesicles or dense core vesicles, suggesting that cilia could receive neurochemical signals at these contacts. These results raise the possibility that cilia might pair with specific partners, possibly using mechanisms analogous to those that control synapse formation. It remains unknown, however, whether the contacts made by neuronal cilia are stereotyped and, if so, what molecular mechanisms pattern them. Here, we show that, in *Caenorhabditis elegans,* neuronal primary cilia attach to specific glial partners using a previously uncharacterized extracellular protein that localizes to cilia-glia contacts. We find that cilia-glia attachments regulate the morphology, calcium dynamics, and molecular compartmentalization of cilia, indicating a critical role in ciliary structure and function. Our results suggest that neuronal cilia not only act as antennae for long-range cellular signals, but also form developmentally programmed contacts that affect how neurons respond to stimuli.

## RESULTS

### Specific neuron-glia attachments develop through a two-step mechanism

Two pairs of *C. elegans* sensory neurons, URX and BAG, provide a powerful model for investigating the development of specific neuron-glia attachments. Each of these neurons extends an unbranched dendrite to the nose tip where it forms a membranous attachment to a protrusion from a specific glial partner (the lateral inner labial socket, here called ILsoL) (Fig. 1A, right)^11–13^. Although the head contains three pairs of ILso glia (dorsal, lateral, and ventral), URX and BAG always attach to ILsoL. The BAG dendrite is positioned near ILsoL in the lateral sensory fascicle, but the URX dendrite lies in the dorsal sensory fascicle and projects across the nose tip to reach ILsoL^12^ (Fig. 1A). The attachments of URX and BAG to ILsoL are conserved in nematodes over ∼100 million years of evolution, suggesting functional importance^14^. Moreover, they develop in a region packed with dozens of glial endings, implying mechanisms for precise neuron-glia pairing.

**Figure 1.**
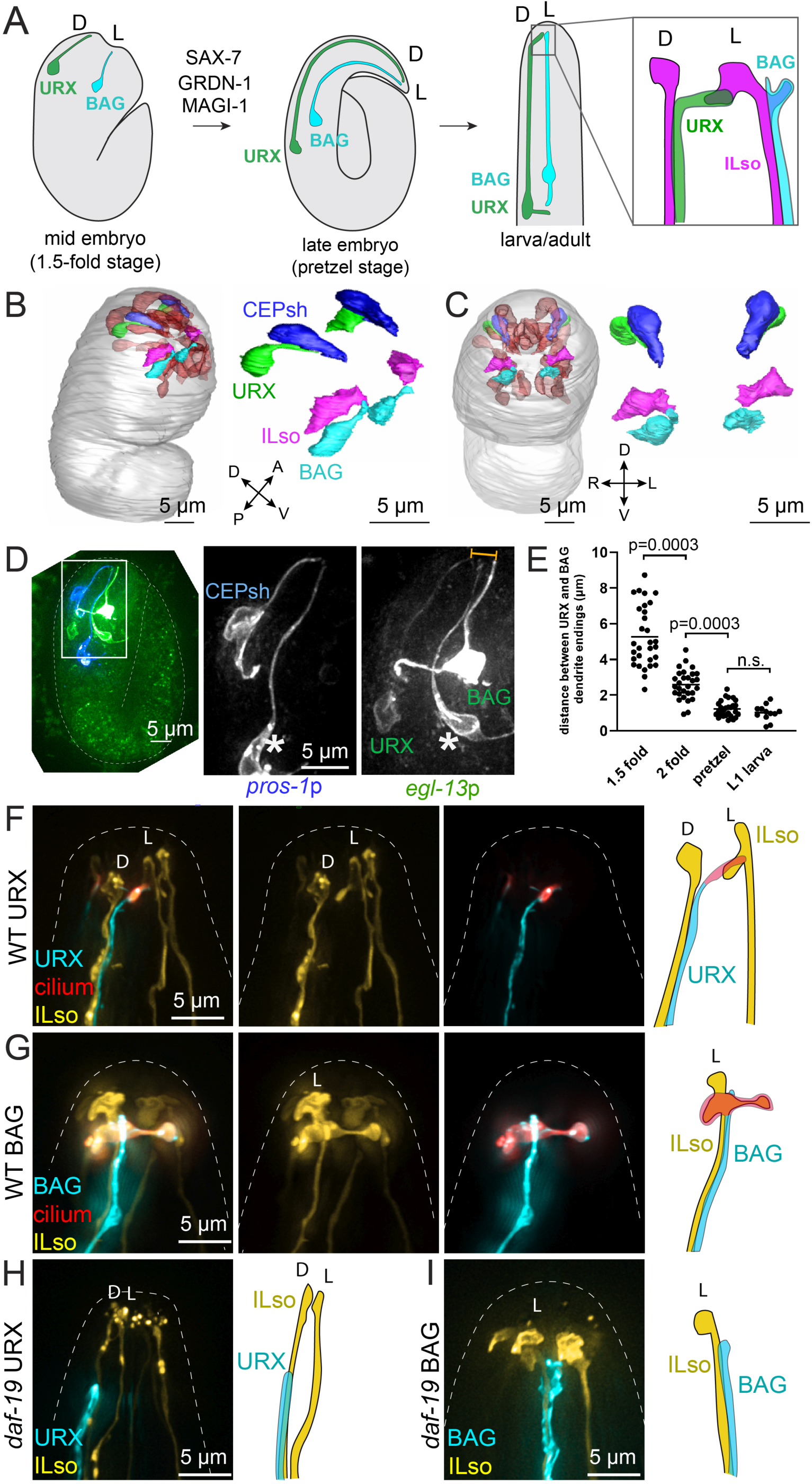
URX and BAG neurons anchor to guidepost glia during embryo elongation, then use primary cilia to attach to their mature glial partner ILsoL. (A) Schematic of URX (green) and BAG (blue) development and attachment to the lateral ILso glia (magenta); D, dorsal; L, lateral. (B, C) Reconstruction of URX (green), BAG (light blue), lateral ILso glia (magenta), dorsal CEPsh glia (dark blue), and other glial cells in the head (brown) from EM dataset of a comma-stage embryo. (D) Maximum-intensity projection of fluorescence in 2-fold embryo showing relative positions of URX, BAG, and CEPsh glia; *egl-13p:myrGFP* labels URX, BAG and AFD (asterisk) neurons in embryos; *pros-1p:myrmCherry* labels several sheath glia in embryos including dorsal CEPsh and amphid sheath (asterisk). Gold brackets mark distance between URX and BAG dendrite endings. (E) Distance between URX and BAG dendrite ending (µm) at various stages of development. Each dot represents an individual URX-BAG pair measurement. P-values determined by Kruskal-Wallis analysis with Dunn’s correction. (F, H) URX dendrite (blue, *flp-8p*), URX cilium (red, *gcy-32p:ARL-13-RFP*), and ILso glia (yellow, *col-53p*) in (F) wild-type and (H) *daf-19(m86)* cilia-defective larvae. (G, I) BAG dendrite (blue, *flp-17p*), BAG cilium (red, *flp-17p:GCY-9-mApple*), and ILso glia (yellow, *col-53p*) in (G) wild-type and (H) *daf-19(m86)* cilia-defective larvae. D, dorsal; L, lateral.

We previously found that URX and BAG dendrites develop through a process called retrograde extension, in which their nascent dendrite endings anchor at the embryonic nose and then stretch up to 10-fold in length during embryo elongation^12,15^. Genetic analysis revealed that anchoring of URX and BAG dendrite endings requires several factors that act in glia, including the scaffolding protein GRDN-1/Girdin (Fig. 1A)^12,15^. If GRDN-1 or other glial factors are disrupted, URX and BAG dendrites detach from the developing nose during embryo elongation, resulting in severely shortened dendrites^12,15^. Thus, URX and BAG dendrites depend on glia in order to extend properly during embryo elongation.

To better understand how neurons interact with glial partners during development, we examined the URX and BAG dendrite endings at the onset of embryo elongation (comma to 1.5-fold stages). We expected that nascent URX and BAG dendrites would anchor to the ILsoL glial cell at this stage.

However, several observations indicated that URX and BAG dendrites initially anchor to different glial cells and only later form their mature attachments to ILsoL glia, implying a two-step mechanism of neuron-glia pairing. First, using electron microscopy of an embryo at the onset of elongation^16,17^, we found that the URX dendrite had not yet exited the dorsal sensory fascicle and its ending was tightly apposed to the dorsal CEPsh glial cell, remaining ∼5 µm away from the ILsoL and BAG endings (Fig. 1B-C, Supp. Fig. S1A). We also observed this arrangement by fluorescence microscopy in 1.5- to 2-fold embryos, the earliest stage at which our cell markers are visible (Fig. 1D). We measured the distance between URX and BAG dendrite endings across embryo development and found that they converge only in post-elongation embryos (pretzel stage) (Fig. 1E). Second, we found that genetic ablation of ILsoL does not affect URX dendrite anchoring (Supp. Fig. S1B-C). Third, we found that expression of GRDN-1 in dorsal CEPsh glia, but not ILsoL, is sufficient for URX dendrite anchoring (Supp. Fig. S1B-C). Together, these results indicate that URX initially attaches to the dorsal CEPsh glial cell, not ILsoL. For BAG, we similarly found that its dendrite ending is closely apposed to a different glial partner (ILsh) during embryo elongation and has not yet wrapped ILsoL (Supp. Fig. S1D), and that ILsoL is not necessary for its dendrite anchoring (Supp. Fig. S1E-F). The lack of embryonic ILsh-specific drivers precluded us from testing if GRDN-1 expression in these cells is sufficient for BAG dendrite anchoring. We concluded that URX and BAG neurons pair with ILsoL glia through a two-step mechanism that involves initial anchoring to other glial partners, prior to formation of the mature attachment (Supp. Fig. S1I). We refer to these intermediate partners as “guidepost glia” to emphasize the similarity to the role of intermediate contacts in axon outgrowth prior to reaching the mature target^18^.

### Primary cilia mediate neuron-glia attachment

To determine how the mature neuron-glia attachment develops, we more closely examined URX and BAG dendrites where they contact ILsoL. URX and BAG have primary (sensory) cilia at their dendrite endings that detect environmental cues (oxygen and carbon dioxide, respectively)^19–22^. We asked whether their primary cilia are involved in forming the neuron-glia attachments. Using fluorescence and electron microscopy, we found that the URX cilium is the only region of the neuron to contact ILsoL (Fig. 1F, Supp. Fig. S1G). As previously described^11–13^, the BAG cilium forms the major contact with ILsoL; however, due to its proximity to ILsoL in the same sensory fascicle, we cannot rule out that other regions of the dendrite may also be attached (Fig. 1G, Supp. Fig. S1H). To test if URX and BAG cilia are required for attachment to ILsoL, we examined *daf-19/*RFX mutants, which do not develop cilia^23,24^. In *daf-19* mutants, the URX dendrite ending fails to exit the dorsal sensory fascicle and does not attach to ILsoL, and BAG dendrites do not wrap ILsoL despite their proximity (Fig. 1H-I). ILsoL also lacks the protrusions around which URX and BAG cilia normally wrap (Fig. 1F-I). Together, these observations suggest that, after dendrite extension is complete, primary cilia outgrowth mediates establishment of the mature neuron-glia attachments.

### BUG-1 is required for cilia-glia attachment

To explore the molecular mechanisms that control cilia-glia attachment, we performed a forward genetic screen (Supp. Fig. S2A-B and Methods) and isolated a mutant, *hmn356*, in which the URX cilium is present but does not attach to ILsoL (Fig. 2A-C). In this mutant, BAG cilia also fail to attach to ILsoL, and ILsoL lacks the protrusions that URX and BAG normally wrap (Fig. 2D-F). Cilia of other sensory neurons remain intact as assessed by gross morphology and dye uptake, indicating that the mutant does not cause general defects in ciliogenesis (Supp. Fig. S2C). We mapped *hmn356* to a nonsense mutation (W85stop) in an uncharacterized gene, *T01D3.1* (Fig. 2G). We used CRISPR/Cas9 to introduce an identical mutation (*hmn404)* or delete the entire locus (*syb9034)* in an otherwise wild-type background (Fig. 2G). Both mutations cause the same phenotype as *hmn356,* demonstrating that *T01D3.1* is necessary for URX and BAG cilia-glia attachment (Fig. 2H). We assigned *T01D3.1* the name *bug-1* for **<u>B</u>**AG and **<u>U</u>**RX **<u>g</u>**lial attachment.

**Figure 2.**
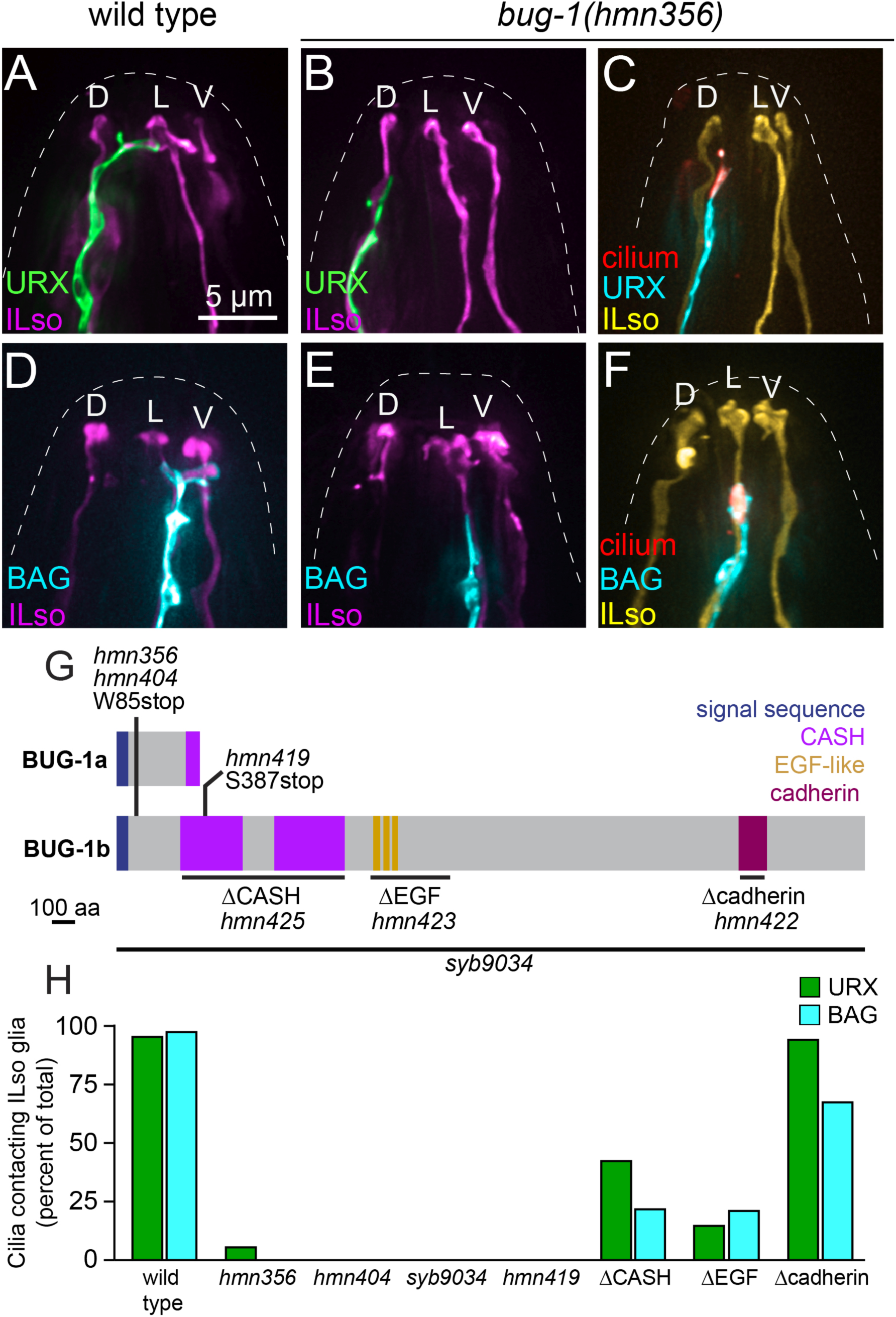
BUG-1 is required for cilia-glia attachment. (A, B) URX (green, *flp-8p*) with ILso glia (magenta, *grl-18p*) in (A) wild-type and (B) *hmn356* larvae. (C) URX cilia in *hmn356* larvae; URX dendrites are blue (*flp-8p*), URX cilia are red (*gcy-32p:ARL-13-RFP*), ILso glia are yellow (*col-53p*). (D, E) BAG (blue, *flp-17p*) with ILso glia (magenta, *grl-18p*) in (D) wild-type and (E) *hmn356* larvae. (F) BAG cilia in *hmn356* larva; BAG dendrites are blue (*flp-17p*), BAG cilia are red (*flp-17p:GCY-9-mApple*), ILso glia are yellow (*col-53p*). (G) Schematic of BUG-1/T01D3.1 showing isoforms, alleles, and domains including signal sequence, CASH (carbohydrate-binding proteins and s̱ ugar ẖydrolases), EGF-like, and cadherin domains. (H) Percent of URX (green) and BAG (blue) cilia contacting the lateral ILso in *bug-1* mutant alleles, n≥50 per genotype.

*bug-1* is predicted to encode a secreted protein with two isoforms: a short form, BUG-1a (39 kDa), and a long form, BUG-1b (352 kDa) with weak similarity to mammalian teneurins, a family of transmembrane adhesion proteins involved in synapse development^25^. To determine which BUG-1 isoform is required for cilia-glia attachment, we used CRISPR/Cas9 to insert a stop codon downstream of the *bug-1a* coding sequence to generate an allele (*hmn419)* that encodes a severely truncated BUG-1b but that does not impact BUG-1a (Fig. 2G). In this mutant, URX and BAG cilia fail to attach to ILsoL, indicating that BUG-1b is necessary for cilia-glia attachment (Fig. 2H). While BUG-1a might have other functions, our results indicate that it is not sufficient for URX and BAG cilia attachment to ILsoL.

BUG-1b contains two CASH domains, several clustered EGF domains, and a cadherin-like domain (Fig. 2G). CASH (<u>ca</u>rbohydrate-binding proteins and <u>s</u>ugar <u>h</u>ydrolases) domains are highly conserved yet virtually unstudied protein domains present across diverse species in >1000 proteins, many with carbohydrate binding functions^26^. We used CRISPR/Cas9 to create in-frame deletions of each set of domains in BUG-1b (Fig. 2G). Deletion of BUG-1b CASH domains (*hmn425)* or clustered EGF-like domains (*hmn423)* caused strong but incompletely penetrant loss of URX and BAG attachments to ILsoL (Fig. 2H). Loss of URX and BAG attachments is usually accompanied by loss of the characteristic ILsoL glial protrusions, but in *bug-1* EGF domain deletion mutants, over half of ILsoL glia retain an aberrantly shaped protrusion – resembling a deflated balloon – that is apposed to, but not wrapped by, the BAG cilium (Supp. Fig. S2D-E). Deletion of the BUG-1b cadherin domain (*hmn422)* had no effect on URX-ILsoL attachment and a modest effect on BAG-ILsoL attachment (Fig. 2H). Together, these results indicate that BUG-1b CASH and EGF domains are critical for cilia-glia attachment and that each domain functions partly independently of the other.

### BUG-1 localizes to the site of cilia-glia attachment

We next sought to determine the localization of BUG-1b. Transcriptomic studies found that *bug-1* mRNA is highly expressed in embryonic and larval URX and BAG neurons, but not in ILso glia^27,28^. Because BUG-1b has an amino-terminal signal peptide and domains associated with cell adhesion, we reasoned that it likely acts at the cell surface or in the extracellular space. We tagged endogenous BUG-1b at its carboxyl-terminus with sfGFP and observed BUG-1b foci that correspond to the positions of URX and BAG cilia (Fig. 3A-C). We also observed BUG-1b foci at every sense organ, each of which consists of one or more ciliated neurons wrapped by a pair of glial cells (amphid, IL, OLQ, OLL, and CEP in the head, Fig. 3A; AQR, ADE, PDE, PQR and phasmid in the body and tail, Supp. Fig. S3A-D). This observation suggests that BUG-1b is present at every neuron-glia attachment in the animal. We did not observe loss of neuron-glia attachments in *bug-1* mutant sense organs (amphid, IL, and CEP; Supp. Fig. S4), although the attachments of some amphid dendrites to the amphid sheath glial cell are mispositioned (Supp. Fig. S4), a finding that will be described in detail elsewhere.

**Figure 3.**
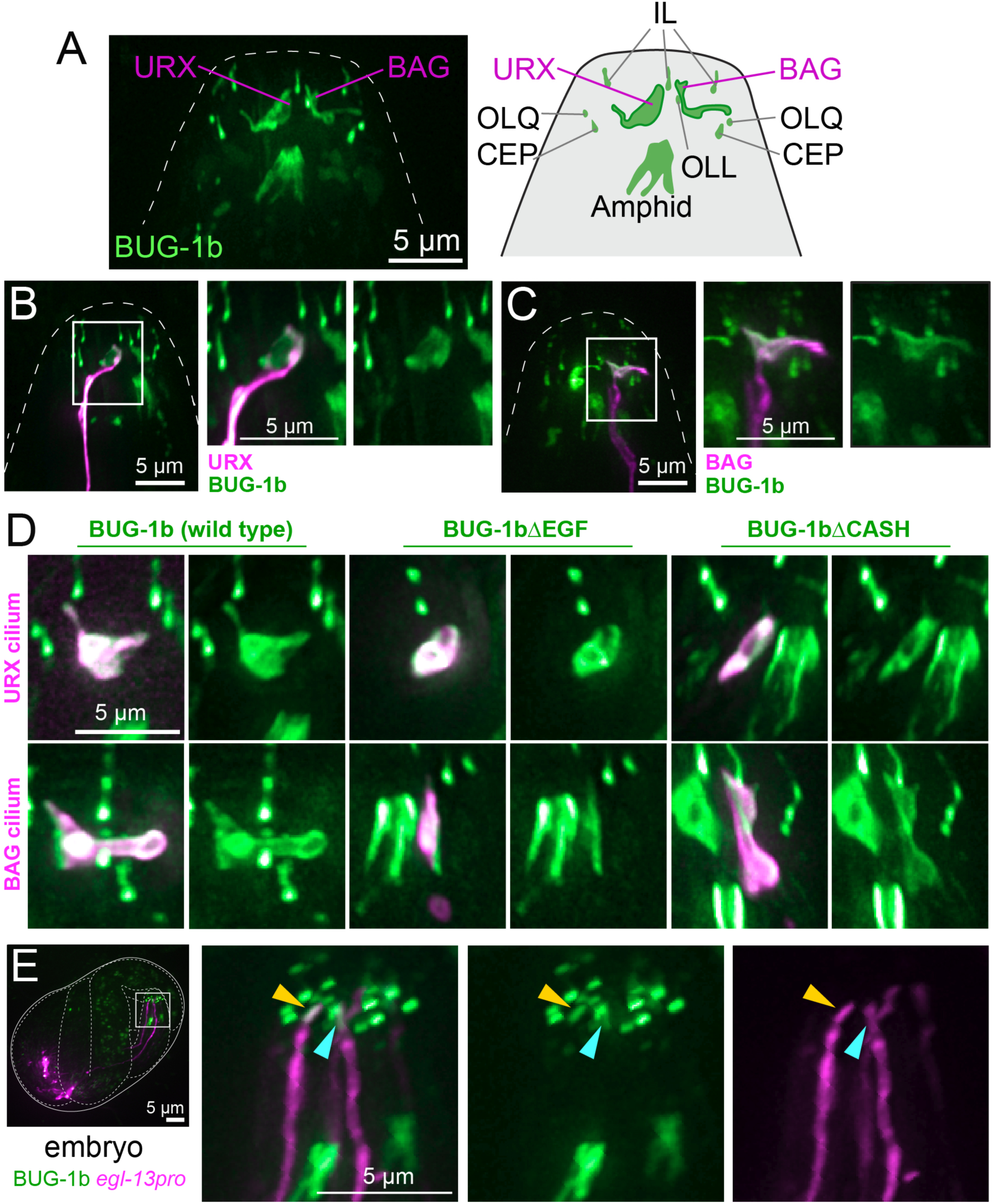
BUG-1b localizes to neuron-glia attachments. (A) Maximum-intensity projection of endogenous BUG-1b-sfGFP localization in the nose of larvae with schematic. (B) BUG-1b-sfGFP with URX neuron (magenta, *flp-8p*). (C) BUG-1b-sfGFP with BAG neuron (magenta, *flp-17p*). (D) BUG-1sfGFP with URX cilium (top) and BAG cilium (bottom) (magenta, *gcy-32p:ARL-13-RFP* and *flp-17p:GCY-9-mApple* respectively) in wild-type, BUG-1bΔEGF, and BUG-1bΔCASH larvae. (E) BUG-1b-sfGFP in late pretzel-stage embryo with URX and BAG neurons (magenta, *egl-13p*) showing localization to developing endings of URX (yellow arrowhead) and BAG (blue arrowhead).

Using URX- and BAG-specific markers, we found that BUG-1b localizes at the sites of attachment to ILsoL, appearing to entirely coat each cilium (Fig. 3B-C). To determine how individual protein domains contribute to BUG-1b localization, we introduced CASH and EGF domain deletions into endogenous BUG-1b tagged with sfGFP. These mutant proteins are properly localized and coat the URX and BAG cilia despite their inability to mediate cilia-glia attachment (Fig. 3D). We conclude that BUG-1b CASH and EGF domains are not required for BUG-1b expression or its localization to cilia, but instead promote attachment to glia.

To determine when BUG-1b is expressed during nervous system development, we examined embryos. BUG-1b expression was first detectable in 2-fold-stage embryos, prior to the outgrowth of URX and BAG cilia (Supp. Fig. S3E). In pretzel-stage embryos, when URX and BAG cilia emerge, BUG-1b localizes to the URX and BAG endings where cilia-glia attachment will occur (Fig. 3E). The localization of BUG-1b to developing and mature URX/BAG-ILsoL contact sites suggests that it plays a direct role in the formation of cilia-glia attachments.

### Cilia-glia attachment is required for activity-dependent URX dendrite remodeling

We took advantage of the *bug-1* mutant to investigate how cilia-glia attachment affects cilia function. URX and BAG neurons sense the respiratory gases oxygen (O_2_) and carbon dioxide (CO_2_), respectively, through guanylate cyclases (GCs) and cyclic GMP-regulated ion channels localized to their cilia^19,21,29,30^. Calcium entry into the cilium depolarizes the neuron, and can also mediate local signaling that is important for sensory adaptation and long-term plasticity^31–37^.

We first asked whether loss of BUG-1b changes how URX neurons respond to O_2_. We did not observe significant differences in URX responses to acute O_2_ stimuli in *bug-1* and wild-type strains (Supp. Fig. S5A-C). Similarly, behavioral responses to acute O_2_ stimuli were not significantly altered in the *bug-1* mutant (Supp. Fig. S5D). Prior work found that, in addition to mediating acute responses to O_2_, URX responds to chronic O_2_ -induced activity over a time scale of days by remodeling its dendrite ending^38^. The URX dendrite ending has a simple morphology in larvae and young (Day 1) adults but, following chronic O_2_-induced activity, it becomes elaborately branched in older (Day 4) adults (Fig. 4). Remodeling of the URX dendrite ending correlates with increased O_2_ avoidance behavior and may be important for an animal to properly navigate natural environments^38^. We observed severely reduced branching of the URX dendrite ending in Day 4 adult *bug-1* mutants compared to wild-type animals (Fig. 4). We also observed reduced branching in animals with genetic ablation of ILsoL, consistent with this defect being caused by failure of cilia to attach to glia (Fig. 4). Together, these results indicate that URX cilia-glia attachment is required for URX responses to chronic, but not acute, sensory stimuli.

**Figure 4.**
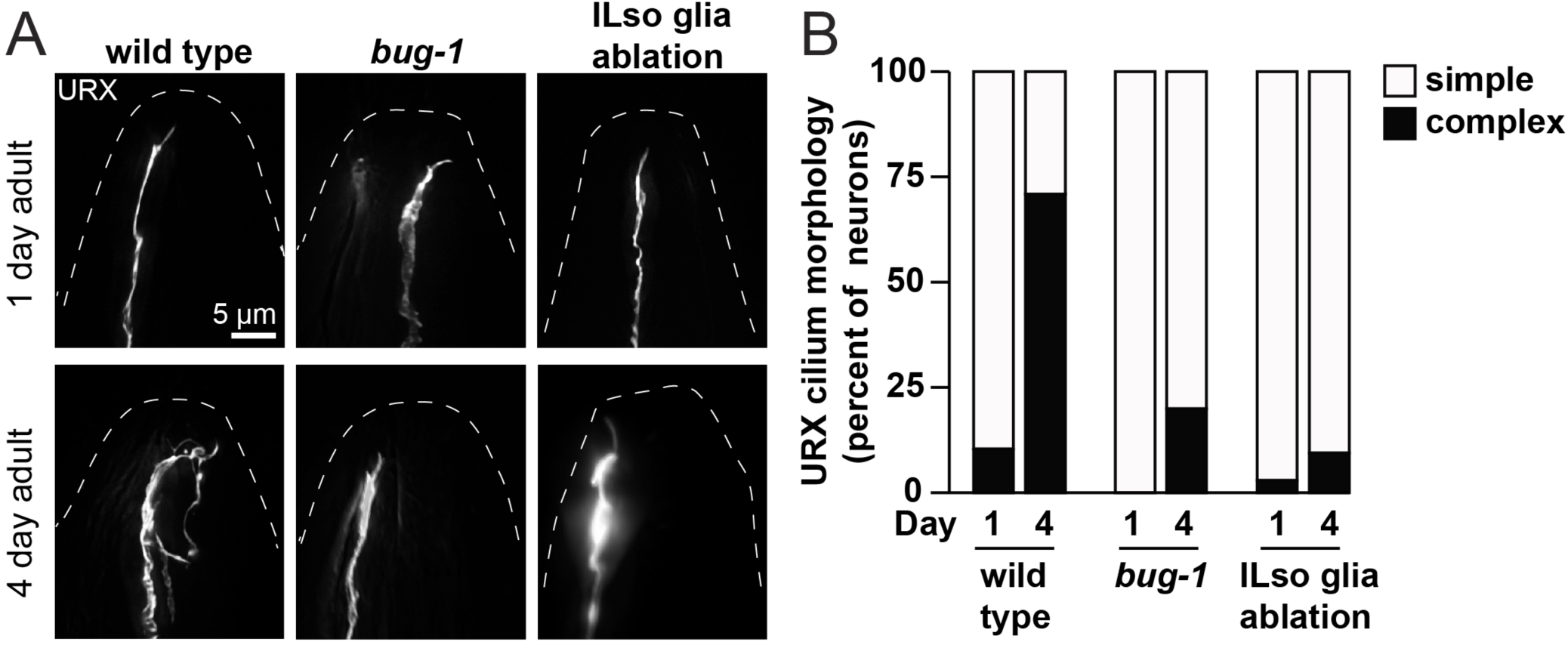
Age- and activity-dependent remodeling of the URX dendrite ending requires glial attachment. (A) URX dendrites (*flp-8p*) in day 1 or 4 of adulthood in wild-type, *bug-1(hmn404),* and ILso ablated (*col-53p:DTA)* animals. (B) Percent of animals with simple or complex URX dendrite endings. Animals with >2 branches on URX dendrite were considered complex. For each group n=28-31.

### Cilia-glia attachment is required for proper calcium dynamics in the BAG cilium

We next examined effects of disrupting cilia-glia attachments on BAG neuron function. We used the fast fluorescent calcium indicator GCaMP6f^39^ to monitor calcium in the cilia and soma of BAG neurons upon CO_2_ exposure (Fig. 5A-B; Supp. Fig. S5E-G). In wild-type animals, ciliary calcium rose quickly, peaked within 500 msec of stimulus onset, and then decreased despite ongoing CO_2_ exposure, a profile that indicates sensory adaptation (Fig. 5B-C,E). Calcium responses in the BAG cilia of *bug-1* mutants were notably different. In *bug-1* mutants, the initial response to stimuli was rapid but then, instead of adapting, ciliary calcium continued to slowly increase, ultimately peaking after the CO_2_ stimulus was removed and several seconds after stimulus onset (Fig. 5B-C, E). The amplitude of the BAG ciliary calcium peak in *bug-1* mutants was slightly but significantly lower than that of wild-type animals (Fig. 5D). These altered cilia calcium dynamics correlated with a modest reduction in avoidance of strong CO_2_ stimuli, but the difference did not reach statistical significance (Supp. Fig. S5H). Importantly, calcium signals in BAG soma appeared normal in *bug-1* mutants, indicating that loss of *bug-1* affects ciliary calcium dynamics and does not generally impair BAG neuron excitability (Supp. Fig. S5E-G). Together, these data suggest that cilia-glia attachments can affect localized calcium signaling, especially in response to sustained stimuli.

**Figure 5.**
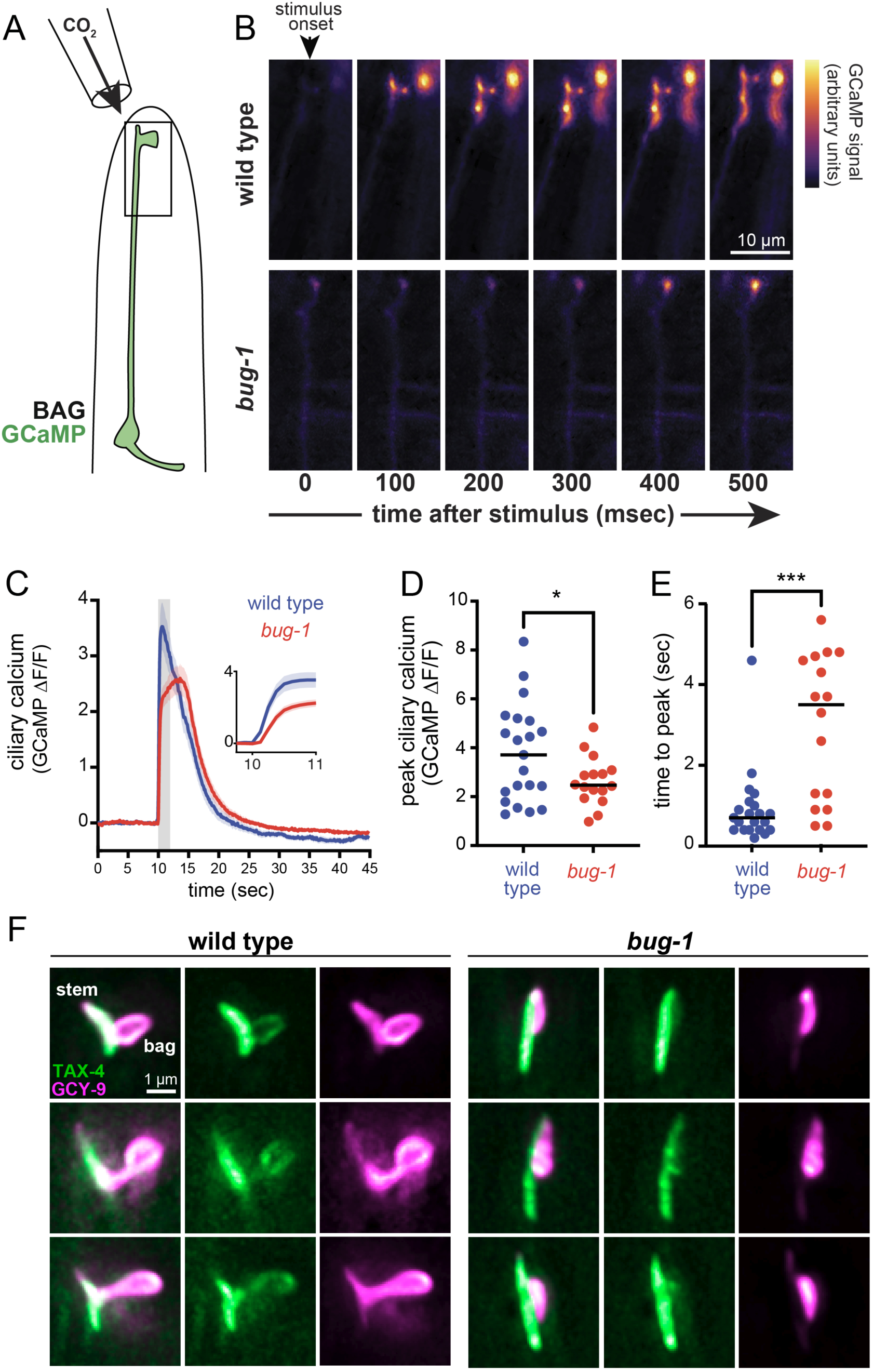
Calcium dynamics in the BAG cilium depend on glial attachment. (A) Schematic of 10% CO_2_ delivery to nose tip of *flp-17*p:GCaMP6 animal glued to coverslip. (B) Representative pseudocolored images showing absolute GCaMP fluorescence in BAG cilia as a time series upon switch to 10% CO_2_ (t=0 msec) in wild-type (top) and *bug-1(hmn404)* (bottom) animals. (C) Quantification of change in GCaMP signal in BAG cilium. Ambient CO_2_ is replaced with 10% CO_2_ from t=10 sec to t=12 sec (gray shading). Inset shows immediate response from t=10 sec to t=11 sec. Solid lines, mean; shaded red or blue, SEM. (D) Peak calcium response and (E) time to peak in sec. *P* values from comparisons of wild-type and *bug-1* responses were computed using Student’s T test with Welch’s correction. * *P* = 0.03, *** *P* = 0.0005*. n =* 21 wild type, *n* = 16 *bug-1*. (F) Maximum-intensity projections of wild-type (left) or *bug-1(hmn404)* (right) BAG cilia expressing endogenous GCY-9-mApple and TAX-4-GFP11 with BAG-specific GFP1-10. Three cilia from different individuals are shown for each genotype.

We hypothesized that these effects might be partly due to altered spatial organization of the BAG cilium upon loss of the cilia-glia attachment. Prior work suggested that the BAG cilium is organized into subcompartments, with the receptor-type guanylyl cyclase GCY-9 and the cyclic nucleotide-gated cation channel subunit TAX-4 preferentially localized to distinct subciliary regions^40^. To test this hypothesis, we visualized endogenous GCY-9 tagged with mApple and endogenous TAX-4 tagged with GFP11, which we combined with BAG-specific expression of GFP1-10. In wild-type animals, we found that TAX-4 was enriched in the stick-like portion of the BAG cilium (“stem” region, Fig. 5F) with weaker localization to the membranous portion of the cilium that wraps the glial protrusion (“bag” region, Fig. 5F). By comparison, GCY-9 localizes to both the stem and bag regions, with higher localization to the bag in some individuals (Fig. 5F). In *bug-1* mutants, these patterns are altered: TAX-4 remains primarily in the stem region, while GCY-9 is localized almost exclusively to the bag region – which, surprisingly, is still present – resulting in severely reduced overlap between the two proteins (Fig. 5F, Supp. Fig. S5I). This change in spatial organization could explain the calcium signaling defects observed in *bug-1* mutant BAG cilia, as the receptor that produces cGMP in response to sensory stimuli is now separated from the cGMP-gated channels that trigger depolarization of the neuron and the calcium influx into the cilium.

## DISCUSSION

The discovery that neuronal cilia make extensive contacts with glia and other cells in the brain has spurred a hunt for molecules that pattern these connections^8–10^. In situ proteomics of mouse brain identified adhesion molecules that are specifically enriched in neuronal cilia^41^, but the complexity of mammalian brain makes it difficult to ascertain their roles. To circumvent this issue, we used the stereotyped anatomy and powerful genetics of *C. elegans* to identify a factor that is essential for cilia-glia attachment. Prior work showed that *C. elegans* glia promote morphogenesis and function of cilia that are ensheathed inside a glial channel or pocket^42–45^, and our results extend this concept to cilia that form peripheral attachments to glia, similar to those in mammalian brain.

Our observation that glial attachments affect how cilia respond to chronic stimuli is reminiscent of roles for glia in promoting changes at synapses in response to repetitive neuron firing. For example, pioneering work showed that perisynaptic Schwann cell glia promote synaptic depression, and subsequent studies revealed diverse mechanisms by which glia – especially astrocytes – modulate synaptic plasticity^46,47^. The genetic tools available in *C. elegans* will enable future work to define the mechanisms by which glial attachments alter ciliary function, including whether glia actively signal to cilia in addition to providing structural support for cilia morphogenesis and compartmentalization.

More broadly, there is evidence that primary cilia attach to neighboring cells in tissues outside the nervous system, suggesting that defining the mechanisms of cilia attachment will have far-reaching implications. For example, primary cilia form long-lasting attachments in liver cholangiocytes *in vivo* and in a kidney-derived cell line *in vitro*^48^. In the pancreas, the primary cilia of insulin-secreting beta cells form cilia-cilia attachments with other beta cells and cilia-cell attachments to other islet cells and to axons that innervate the organ^48,49^. These attachments are ideally positioned to regulate ciliary signaling. Our discovery of genetically specified cilia-glia attachments in *C. elegans* raises the intriguing possibility that patterned cilia-cell attachments that modulate ciliary signaling may be an evolutionary ancient feature of animal tissues with widespread impacts on organismal physiology.

## METHODS

### Strains and plasmids

All strains were made in the N2 background unless otherwise stated. Animals were grown at 20°C on nematode growth media (NGM) plates seeded with *E. coli* OP50 bacteria^50^. Transgenic strains were generated using standard techniques^51^ with injections of 100-160 ng/µl DNA (5-100 ng/µl per plasmid).

### Electron microscopy

Analysis was performed on the dataset “c_elegans_embryo_345min” obtained by FIB-SEM imaging of a 27.5 µm × 26.5 µm × 41.0 µm volume (1641 sections each of approximately 25-nm thickness) as previously described^16,17^. Cells were manually traced using IMOD as individual objects using the Model mode. All three-dimensional graphical models were generated from these tracings in IMOD using the Model View mode^52^.

### Fluorescence microscopy and image processing

Larval and adult animals were mounted on 2% agarose pads containing 50 mM sodium azide. Embryos were left in a drop of sodium azide for ∼2 min until twitching ceased and were then mounted on 2% agarose pads. Image stacks were collected on a DeltaVision Core imaging system (Applied Precision) with UApo 40x/1.35 NA, PlanApo 60x/1.42 NA, and UPlanSApo 100x/1.40 NA oil immersion objectives and a CoolSnap HQ2 camera. Images were deconvolved using Softworx 5.5 (Applied Precision). Maximum intensity projections were generated in ImageJ (Fiji). The brightness and contrast of each projection were linearly adjusted, fluorescent signals were pseudo-colored, and merged images were generated using the Screen layer mode in Affinity Photo 1.10.1.

### Quantification of dendrite lengths and distance between URX and BAG dendrite endings

Dendrite lengths were measured in L3-L4 stage animals using Softworx 5.5. Dendrite length was measured from the point at which the dendrite connects with the cell body to its distal terminus at the nose. To normalize for variance in animal size and cell body position, the distance from the cell body to the tip of the dendrite was divided by the distance from the cell body to the tip of the nose. The distance between URX and BAG dendrite endings in embryos and L1 larvae was measured from optical stacks acquired with 0.2 µm between optical sections. Statistical analysis was performed using GraphPad Prism 9 software.

### Forward genetic screen and mapping of mutations

In order to identify mutants with altered positioning of the URX dendrite ending, a *mapk-15* mutant background was used to increase the length of the URX dendrite ending^53^ and thus facilitate visual screening (Supp. Fig. S2B). *mapk-15(hmn5); ynIs78[flp-8p:GFP]; hmnIs47[grl-18p:mCherry]* animals at the L4 stage were mutagenized using 70 mM ethyl methanesulfonate (EMS, Sigma) at 22°C for 4 h. F2 progeny were screened using a fluorescence stereomicroscope, and animals with abnormal URX dendrite position were isolated to individual plates. The *mapk-15(hmn5)* mutation was crossed out of individual mutant strains by selecting animals that lack URX overgrowth but exhibit altered URX cilia position assessed using a UPlanSApo 100×/1.40 NA oil immersion objective.

Using linkage mapping and SNP analysis, *hmn356* was mapped to the interval between 9 Mb and 19 Mb on chromosome V. Following whole-genome sequencing to identify candidate mutations in this interval, *hmn356* was crossed to a strain bearing the visible flanking markers *dpy-11 (*6.5 Mb; 0 cM) and *unc-76* (15 Mb; 7.3 cM). F2 recombinants in this region were identified by selecting Dpy nonUnc (n=15) or Unc nonDpy (n=4) isolates. These isolates were assessed for the *hmn356* phenotype (altered URX cilia position, using a UPlanSApo 100×/1.40 NA oil immersion objective) and the presence or absence of candidate mutations by PCR. This approach led to the identification of a small linkage interval that contained a W85stop mutation in the previously uncharacterized gene *T01D3.1*, which we named *bug-1*.

### Generation of alleles by genome editing

The CRISPOR web tool^54^ was used to identify potential guide RNA sequences near target regions.

The *bug-1(syb9034)* and *tax-4(syb10450)* alleles were generated by SunyBiotech (Fuzhou, China).

### Calcium imaging of URX and BAG neurons

Young adults carrying a GCaMP transgene expressed in URX or BAG neurons were immobilized with surgical glue (SurgiLock) on a 2% agarose pad. Animals were transferred to an inverted microscope (Nikon Eclipse Ti) equipped with an LED light engine (CoolLED) and high-sensitivity CMOS cameras (Hamamatsu Orca Fusion-BT). Image series were acquired with a 20x long working distance air objective (NA 0.8) at 10 frames/sec for CO_2_ stimuli and 2 frames/sec for O_2_ stimuli. Stimuli were delivered to immobilized animals through glass pipettes that superfused immobilized animals with humidified gas mixtures (AirGas). Flow of gases through local-perfusion pipettes was controlled by solenoid valves (NResearch) using custom-built valve controllers. Image acquisition and valve control was coordinated by a Virtex controller and VisiView software (VisiTron Systems). For each recording, background fluorescence was subtracted and GCaMP fluorescence in regions of interest was normalized to the mean fluorescence from the first 100 frames of the video, i.e. before stimulus presentation. Image analysis was performed using Fiji. Peak response and time to peak response for each recording was determined by identifying the maximum GCaMP signal in the recording after stimulus presentation and the associated time.

### Measuring URX- and BAG-dependent behaviors

BAG-dependent CO_2_ avoidance and URX-dependent O_2_ response behaviors were analyzed as previously described^55^. Briefly, avoidance of CO_2_ by wild-type and *bug-1* mutant animals was tested in custom-made circular arenas with 6 cm diameter. Inlets for control air and air with 10% CO_2_ were place on opposite sides of the chamber 1 cm from the midline, and gas mixes were pushed at 1.5 ml/min using a two-channel syringe pump (New Era). Approximately 50 animals were transferred into the arena and positioned at the midline. After 20 min, the number of animals on the control and high CO_2_ sides of the arena were counted and an avoidance index (AI) was calculate as follows: AI = [number on air side - number on CO_2_ side] / [number on air side + number on CO_2_ side]. O_2_ response behaviors were measured in the same arenas. For these assays, we introduced an *npr-1* mutation that renders animals highly responsive to activation of URX neurons by O_2_^56^. *npr-1* and *bug-1; npr-1* mutants were adapted to 7% O_2_ for 5 min. We acquired videos of animals at 1 frame/sec as the atmosphere in the chamber was switched from 7% O_2_ to 21% O_2_. Atmosphere was continuously pushed into the chamber at 1.5 ml/min using a two-channel syringe pump (New Era). Switching between low and high O_2_ atmospheres was controlled by a solenoid shuttle valve (NResearch).

Videos were analyzed by a multi-worm tracker^57^ and O_2_ responses were measured as changes to the instantaneous speed of animals in the chamber.

## ACKNOWLEDGMENTS

We thank WormBase and the CGC, which is funded by NIH Office of Research Infrastructure

Programs (P40 OD010440).

