## Supplementary Data for "A stereotyped glial attachment determines the morphology and function of neuronal cilia"

**SUPPORTING TABLE S1**

**Strains generated for this study**

| Strain | Genotype | Figure |
|---|---|---|
| CHB4205 | *hmnEx2294[pros-1p:myrmCherry + egl-13p:myrGFP + pRF4]* | 1D, 1E |
| CHB4547 | *hmnEx2444[col-53p:myrYFP + flp-8p:CFP + gcy-32p:ARL-13-TAG-RFP + pRF4]* | 1F |
| CHB5785 | *hmnEx2559[col-53p:myrYFP + flp-17p:CFP + flp-17p:GCY-9-mApple + pRF4]* | 1G |
| CHB4649 | *daf-19(ma86); daf-12(sa204); hmnEx2444[col-53p:myrYFP + flp-8p:CFP + gcy-32p:ARL-13-TAG-RFP + pRF4]* | 1H |
| CHB5796 | *daf-19(ma86); daf-12(sa204); hmnEx2559[col-53p:myrYFP + flp-17p:CFP + flp-17p:GCY-9-mApple + pRF4]* | 1I |
| CHB4798 | *ynIs78[flp-8p:GFP]; hmnIs47[grl-18p:mApple]; hmnEx2529[col-53p:DT-A + pRF4]* | S1C, 4A, 4B |
| CHB4441 | *grdn-1(ns303); ynIs78[flp-8p:GFP]; hmnEx2379[col-53p:GRDN-1a + pRF4]* | S1C |
| CHB4405 | *grdn-1(ns303); ynIs78[flp-8p:GFP]; hmnEx2377[K08D12.4p:GRDN-1a + pRF4]* | S1C |
| CHB5784 | *oyIs82[flp-17p:GFP + unc-122p:dsRed]; hmnIs47[grl-18p:mApple]; hmnEx2529[col-53p:DT-A + pRF4]* | S1F |
| CHB4406 | *grdn-1(ns303); oyIs82[flp-17p:GFP + unc-122p:dsRed]; hmnEx2378[col-53p:GRDN-1a + pRF4]* | S1F |
| CHB5718 | *grdn-1(ns303); oyIs82[flp-17p:GFP + unc-122p:dsRed]; hmnEx2377[K08D12.4p:GRDN-1a + pRF4]* | S1F |
| CHB4923 | *bug-1(hmn356); ynIs78[flp-8p:GFP]; hmnIs47[grl-18p:mApple]* | 2B |
| CHB5819 | *bug-1(hmn356); hmnEx2444[col-53p:myrYFP + flp-8p:CFP + gcy-32p:ARL-13-TAG-RFP + pRF4]* | 2C |
| CHB5025 | *oyIs82[flp-17p:GF + unc-122pro:dsRed]; hmnIs47[grl-18p:mApple]* | 2D, 2H, S2D, S2E |
| CHB4791 | *bug-1(hmn356); oyIs82[flp-17p:GFP + unc-122p:dsRed]; hmnIs47[grl-18p:mApple]* | 2E, 2H |
| CHB4916 | *bug-1(hmn356); hmnEx2559[col-53p:myrYFP + flp-17p:CFP + flp-17p:GCY-9-mApple + pRF4]* | 2F |
| CHB4799 | *bug-1(hmn356); hmnEx1575[flp-8p:mApple + flp-8p:myrmApple + grl-18p:GFP + grl-18p:myrGFP + pRF4]* | 2H |
| CHB5045 | *bug-1(hmn404); hmnEx1575[flp-8p:mApple + flp-8p:myrmApple + grl-18p:GFP + grl-18p:myrGFP + pRF4]* | 2H, 4A, 4B |
| CHB5009 | *bug-1(hmn404); oyIs82[flp-17p:GFP+ unc-122p:dsRed]; hmnIs47[grl-18p:mApple]* | 2H, S2D, S2E |
| CHB5797 | *bug-1(syb9034); hmnEx1575[flp-8p:mApple + flp-8p:myrmApple + grl-18p:GFP + grl-18p:myrGFP + pRF4]* | 2H |
| CHB5072 | *bug-1(syb9034); oyIs82[flp-17p:GFP + unc-122p:dsRed]; hmnIs47[grl-18p:mApple]* | 2H |
| CHB5798 | *bug-1(hmn419); hmnEx1575[flp-8p:mApple + flp-8p:myrmApple + grl-18p:GFP + grl-18p:myrGFP + pRF4]* | 2H |
| CHB5104 | *bug-1(hmn419); oyIs82[flp-17p:GFP + unc-122p:dsRed];* | 2H |

| | | |
|---|---|---|
| | *hmnIs47[grl-18p:mApple]* | |
| CHB5799 | *bug-1(hmn425); hmnEx1575[flp-8p:mApple + flp-8p:myrmApple + grl-18p:GFP + grl-18p:myrGFP + pRF4]* | 2H |
| CHB5160 | *bug-1(hmn425); oyIs82[flp-17p:GFP + unc-122p:dsRed]; hmnIs47[grl-18p:mApple]* | 2H |
| CHB5186 | *bug-1(hmn423); hmnEx1575[flp-8p:mApple + flp-8p:myrmApple + grl-18p:GFP + grl-18p:myrGFP + pRF4]* | 2H |
| CHB5130 | *bug-1(hmn423); oyIs82[flp-17p:GFP + unc-122p:dsRed]; hmnIs47[grl-18pro:mApple]* | 2H, S2D, S2E |
| CHB5185 | *bug-1(hmn422); hmnEx1575[flp-8p:mApple + flp-8p:myrmApple + grl-18p:GFP + grl-18p:myrGFP + pRF4]* | 2H |
| CHB5129 | *bug-1(hmn422); oyIs82[flp-17p:GFP + unc-122p:dsRed]; hmnIs47[grl-18pro:mApple]* | 2H |
| CHB6431 | *mapk-15(hmn5); ynIs78[flp-8p:GFP]; hmnIs47[grl-18p:mApple]* | S2B |
| CHB4737 | *bug-1(hmn356); mapk-15(hmn5); ynIs78[flp-8p:GFP]; hmnIs47[grl-18p:mApple]* | S2B |
| CHB5782 | *bug-1(hmn404)* | S2C, S5H |
| CHB5423 | *hmn467[BUG-1-sfGFP]* | 3A, S3A, S3B |
| CHB5546 | *hmn467[BUG-1-sfGFP]; hmnEx2757[flp-8p:mApple + flp-8p:myrmApple + pRF4]* | 3B |
| CHB5547 | *hmn467[BUG-1-sfGFP]; hmnEx2758[flp-17p:mApple + flp-17p:myrmApple + pRF4]* | 3C |
| CHB5633 | *hmn467[BUG-1-sfGFP]; hmnEx2779[gcy-32p:ARL-13-TAG-RFP + pRF4]* | 3D |
| CHB5632 | *hmn467[BUG-1-sfGFP]; hmnEx2778[flp-17p:GCY-9-mApple + pRF4]* | 3D |
| CHB5786 | *hmn479[BUG-1ΔEGF-sfGFP]; hmnEx2779[gcy-32p:ARL-13-TAG-RFP + pRF4]* | 3D |
| CHB5645 | *hmn479[BUG-1ΔEGF-sfGFP]; hmnEx2778[flp-17p:GCY-9-mApple + pRF4]* | 3D |
| CHB5787 | *hmn480[BUG-1ΔCASH-sfGFP]; hmnEx2779[gcy-32p:ARL-13-TAG-RFP, pRF4]* | 3D |
| CHB5783 | *hmn480[BUG-1ΔCASH-sfGFP]; hmnEx2778[flp-17pro:GCY-9-mApple, pRF4]* | 3D |
| CHB5471 | *hmn467[BUG-1-sfGFP]; hmnEx2729[egl-13p:mCherry + egl-13p:myrmCherry]* | 3E, S3E |
| CHB5800 | *hmn467[BUG-1-sfGFP]; hmnIs123[dat-1p:mApple]* | S3C, S3D |
| CHB5801 | *hmnIs123[dat-1p:mApple] ; irIs67[hlh-17p:GFP + unc-119(+)]* | S4A |
| CHB5802 | *bug-1(hmn404); hmnIs123[dat-1p:mApple] ; irIs67[hlh-17p:GFP + unc-119(+)]* | S4A |
| CHB5788 | *hmnIs47[grl-18p:mApple]; hmnEx1948[flp-3p:YFP + pRF4]* | S4B |
| CHB5789 | *bug-1(hmn404); hmnIs47[grl-18p:mApple]; hmnEx1948[flp-3p:YFP + pRF4]* | S4B |
| CHB5637 | *ksIs2[daf-7p:GFP]; hmnEx2780[F16F9.3p:mApple + pRF4]* | S4C |
| CHB5638 | *bug-1(hmn404); ksIs2[daf-7p:GFP]; hmnEx2780[F16F9.3p:mApple + pRF4]* | S4C |
| FQ845 | *wzEx165*[*flp-17p:GCaMP6f + unc-122p:mCherry*] | 5B-E, S5E-G |
| FQ3280 | *bug-1(hmn404); wzEx165[flp-17p:GCaMP6f + unc-* | 5B-E, S5E-G |

| | | |
|---|---|---|
| | *122p:mCherry]* | |
| CHB5803 | *gcy-9(hmn497)[gcy-9-mApple]; tax-4(syb10450)[tax-4-GFP11]; hmnEx2821[flp-17p:GFP1-10 + unc-122p:RFP]* | 5F, S5I |
| CHB5804 | *bug(hmn404); gcy-9(hmn497)[gcy-9-mApple]; tax-4(syb10450)[tax-4-GFP11]; hmnEx2821[flp-17p:GFP1-10 + unc-122p:RFP]* | 5F, S5I |
| CHB4835 | *hmnIs121[flp-8p:GCaMP6f + unc-122p:RFP]* | S5A-C |
| CHB5562 | *bug-1(hmn404); hmnIs121[flp-8p:GcamP6 + unc-122p:RFP]* | S5A-C |
| CHB5154 | *bug-1(hmn404); npr-1(ad609)* | S5D |

**SUPPORTING TABLE S2**

**Other strains used in this study**

| **Strain** | **Genotype** | **Figure** | **Source** |
|---|---|---|---|
| NY2078 | *ynIs78[flp-8p:GFP]* | S1B, S1C | Kim and Li, 2004 |
| CHB1402 | *grdn-1(ns303); ynIs78[flp-8p:GFP]* | S1B, S1C | Cebul et al., 2020 |
| CHB2231 | *grdn-1(ns303); ynIs78[flp-8p:GFP]; hmnEx696[pros-1p:GRDN-1a + pRF4]* | S1C | Cebul et al., 2020 |
| PY8503 | *oyIs82[flp-17p:GFP + unc-122pro:dsRed]* | S1E, S1F | Astrid Cornils and Piali Sengupta |
| CHB1409 | *grdn-1(ns303); oyIs82[flp-17p:GFP + unc-122p:dsRed]* | S1E, S1F | Cebul et al., 2020 |
| CHB1393 | *grdn-1(ns303); oyIs82[flp-17p:GFP + unc-122p:dsRed]; hmnEx696[pros-1p:GRDN-1a + pRF4]* | S1F | Cebul et al., 2020 |
| CHB3310 | *ynIs78[flp-8p:GFP]; hmnIs47[grl-18p:mApple]* | 2A, 4A, 4B | Karolina Mizeracka |
| DA609 | *npr-1(ad609)* | S5D | de Bono and Bargmann, 1998 |

**SUPPORTING TABLE S3**

**Plasmids generated for this study**

| **Plasmid** | **Description** |
|---|---|
| pLW4 | *egl-13p:myrGFP* |
| pLW7 | *pros-1p:myrmCherry* |
| pLW122 | *col-53p:DT-A* |
| pLW41 | *col-53p:GRDN-1a* |
| pLW42 | *K08D12.4p:GRDN-1a* |
| pLW12 | *col-53p:myrYFP* |
| pLW11 | *egl-13p:myrmCherry* |
| pMH424 | *flp-3p:YFP* |
| pLW115 | *flp-8p:GCaMP6f* |
| pLW136 | *flp-17p:GFP1-10* |

**SUPPORTING TABLE S4**

**Alleles generated in this study**

| **Allele** | **Sequence** | **Notes** |
|---|---|---|
| *bug-1(hmn356)* | ACACAGAATGCCAAAAGGACATTG[G>A]AAAGTCGCTCATTACACAAATGCAA | W85STOP mutation isolated from EMS mutagenesis screen |
| *bug-1(hmn404)* | See *bug-1(hmn356)* | Missense mutation from *bug-1(hmn356*) introduced into wild-type background by CRISPR/Cas9 editing |
| *bug-1(syb9034)* | gaagagtcgaatagtatgtgacgcg[13,338 bp deletion]tttgaatttgcgggtttatctattg | deletion spanning entire endogenous *bug-1* locus |
| *bug-1(hmn419)* | GAAGTTCATCTTATT[C>A]GAACAAGGCTACTGG | S387STOP mutation, disrupts BUG-1b but not BUG-1a |
| *bug-1(hmn425)* | GATGGAGGGGAATTGAGTTCCGTAA[3,686 bp deletion]AGGAATTTATACAGTTACAGTGGAC | ΔCASH in-frame deletion, spanning CASH domains of endogenous *bug-1* locus |
| *bug-1(hmn423)* | TGCGCTGACTACTGTAGTTCCTGTT[1,119 bp deletion]ATTGACAAACGATTCCTGGAACATC | ΔEGF in-frame deletion, spanning EGF-like domains of endogenous *bug-1* locus |
| *bug-1(hmn422)* | CAACTCCAGAAGATGAGAACCCACA[377 bp deletion]CTTCGGAGATTATATTCCAGCTCCG | Δcadherin in-frame deletion, spanning cadherin domain of endogenous *bug-1* locus |
| *bug-1(hmn467)* | GGAGTGTGCTTCAACCACTGC[C>A]GCG[ATGAGCAAAGGAGAAGAACTTTTCACTGGAGTTGTCCCAATTCTTGTTGAATTAGATGGTGATGTTAATGGGCACAAATTTTCTGTCCGTGGAGAGGGTGAAGGTGATGCTACAAACGGAAAACTCACCCTTAAATTTATTTGCACTACTGGAAAACTACCTGTTCCGTGGCCAACACTTGTCACTACTCTGACCTATGGTGTTCAATGCTTTTCCCGTTATCCGGATCACATGAAACGGCATGACTTTTTCAAGAGTGCCATGCCCGAAGGTTATGTACAGGAACGCACTATATCTTTCAAAGATGACGGGACCTACAAGACGCGTGCTGAAGTCAAGTTTGAAGGTGATACCCTTGTTAATCGTATCGAGTTAAAGGGTATTGATTTTAAAGAAGATGGAAACATTCTTGGACACAAACTCGAGTACAACTTTAACTCACACAATGTATACATCACGGCAGACAAACAAAAGAATGGAATCAAAGCTAACTTCAAAATTCGCCACAACGTTGAAGATGGTTCCGTTCAACTAGCAGACCATTATCAACAAAATACTCCAATTGGCGATGGCCCTGTCCTTTTACCAGACAACCATTACCTGTCGACACAATCTGTCCTTTCGAAAGATCCCAACGAAAAGCGTGACCACATGGTCCTTCTTGAGTTTGTAACTGCTGCTGGGATTACACATGGCAT | sfGFP insertion near carboxyl terminus of endogenous *bug-1* locus and silent mutation to disrupt PAM site |

| | | |
|---|---|---|
| | GGATGAGCTCTACAAA]TTATCTGCATATTCGAAAATCATAT | |
| *bug-1(hmn467hmn479)* | See *hmn467* and *hmn423* | ΔEGF deletion introduced into *hmn467* background |
| *bug-1(hmn467hmn480)* | See *hmn467* and *hmn425* | ΔCASH deletion introduced into *hmn467* background |
| *gcy-9(hmn497)* | CCTTGAAGGAAGAACCGGCAAACAA[ATGGTGAGCAAGGGCGAGGAGAATAACATGGCCATCATCAAGGAGTTCATGCGCTTCAAGGTGCACATGGAGGGCTCCGTGAACGGCCACGAGTTCGAGATCGAGGGCGAGGGCGAGGGCCGCCCCTACGAGGCCTTTCAGACCGCTAAGCTGAAGGTGACCAAGGGTGGCCCCCTGCCCTTCGCCTGGGACATCCTGTCCCCTCAGTTCATGTACGGCTCCAAGGTCTACATTAAGCACCCAGCCGACATCCCCGACTACTTCAAGCTGTCCTTCCCCGAGGGCTTCAGGTGGGAGCGCGTGATGAACTTCGAGGACGGCGGCATTATTCACGTTAACCAGGACTCCTCCCTGCAGGACGGCGTGTTCATCTACAAGGTGAAGCTGCGCGGCACCAACTTCCCCTCCGACGGCCCCGTAATGCAGAAGAAGACCATGGGCTGGGAGGCCTCCGAGGAGCGGATGTACCCCGAGGACGGCGCCCTGAAGAGCGAGATCAAGAAGAGGCTGAAGCTGAAGGACGGCGGCCACTACGCCGCCGAGGTCAAGACCACCTACAAGGCCAAGAAGCCCGTGCAGCTGCCCGGCGCCTACATCGTCGACATCAAGTTGGACATCGTGTCCCACAACGAGGACTACACCATCGTGGAACAGTACGAACGCGCCGAGGGCCGCCACTCCACCGGCGGCATGGACGAGCTGTACAAGGAATTCAAC]TGAgattgatttggattcaaagatt | GCY-9-mApple |
| *tax-4(syb10450)* | AACTGAATCTGAATCCTTGCTCAAA[GTACCGGTAGAAAAACGTGACCACATGGTCCTTCATGAGTATGTAAATGCTGCTGGGATTACAGGTGGCTCTGGAGGTAGAGATCATATGGTTCTCCACGAATACGTTAACGCCGCAGGCATCACTGGCGGTAGTGGAGGACGCGACCATATGGTACTACATGAATATGTCAATGCAGCCGGAATAACCGGAGGGTCCGGAGGCCGGGATCACATGGTGCTGCATGAGTATGTGAACGCGGCGGGTATAACTGGTGGGTCGGGCGGACGAGACCATATGGTGCTTCACGAATACGTAAACGCAGCTGGCATTACTGGCGGATCAGGTGGCAGGGATCACATGGTACTCCATGAGTACGTGAACGCTGCTGGAATCACAGGCGGTAGCGGCGGTCGGGACCATATGGTCCTGCACGAATATGTCAATGCTGCCGGTATCACC] TAGttatatttttaatttttaact | TAX-4-7XGFP11 |

**SUPPORTING TABLE S5**

**Guide RNAs used for CRISPR/Cas9-mediated genome editing**

| Oligonucleotide | Sequence | Notes |
|---|---|---|
| bug-1-sgRNA W85stop | CAGAATGCCAAAAAGGACAT | W85stop mutation in *bug-1* |
| bug-1-sgRNA S387stop | AAGTTCATCTTATTCGAACA | S387stop mutation in *bug-1* |
| bug-1-sgRNA Δbug-1 locus #1 | AGTCGAATAGTATGTGACGC | 5' end of *bug-1* locus to generate Δbug-1 strain |
| bug-1-sgRNA Δbug-1 locus #2 | TTTCAAAGATTTGAATTTGC | 3' end of *bug-1* locus to generate Δbug-1 strain |
| bug-1-sgRNA ΔCASH #1 | GAGATGATTCTGATGAGTTA | 5' end of CASH domains to generate ΔCASH deletion |
| bug-1-sgRNA ΔCASH #2 | AACAATTTCACAAATAACGG | 3' end of CASH domains to generate ΔCASH deletion |
| bug-1-sgRNA ΔEGF #1 | AAAGAATCTGGGACTGTAAC | 5' end of EGF-like domains to generate ΔEGF deletion |
| bug-1-sgRNA ΔEGF #2 | AGGAATCGTTTGTCAATTGT | 3' end of EGF-like domains to generate ΔEGF deletion |
| bug-1-sgRNA Δcadherin #1 | ACTGAAGTTGGTCAAACTGT | 5' end of cadherin domain to generate Δcadherin deletion |
| bug-1-sgRNA Δcadherin #2 | CTGACACCAATGATTCACTT | 3' end of cadherin domain to generate Δcadherin deletion |
| bug-1-sgRNA sfGFP | TTTCGAATATGCAGATAACG | for insertion of sfGFP near *bug-1* carboxyl terminus |
| gcy-9-sgRNA mApple | GGCAAACAATGAGATTGATT | for insertion of mApple at *gcy-9* carboxyl terminus |
| tax-4-sgRNA GFP11 #1 | TTTGTTTGTCGGAGACGTGG | for insertion of GFP11 at *tax-4* carboxyl terminus |
| tax-4-sgRNA GFP11 #2 | GATTCAGATTCAGTTCCCGT | for insertion of GFP11 at *tax-4* carboxyl terminus |
| tax-4-sgRNA GFP11 #3 | AAAATATAACTATTTGAGCA | for insertion of GFP11 at *tax-4* carboxyl terminus |

SUPPORTING FIGURE S1

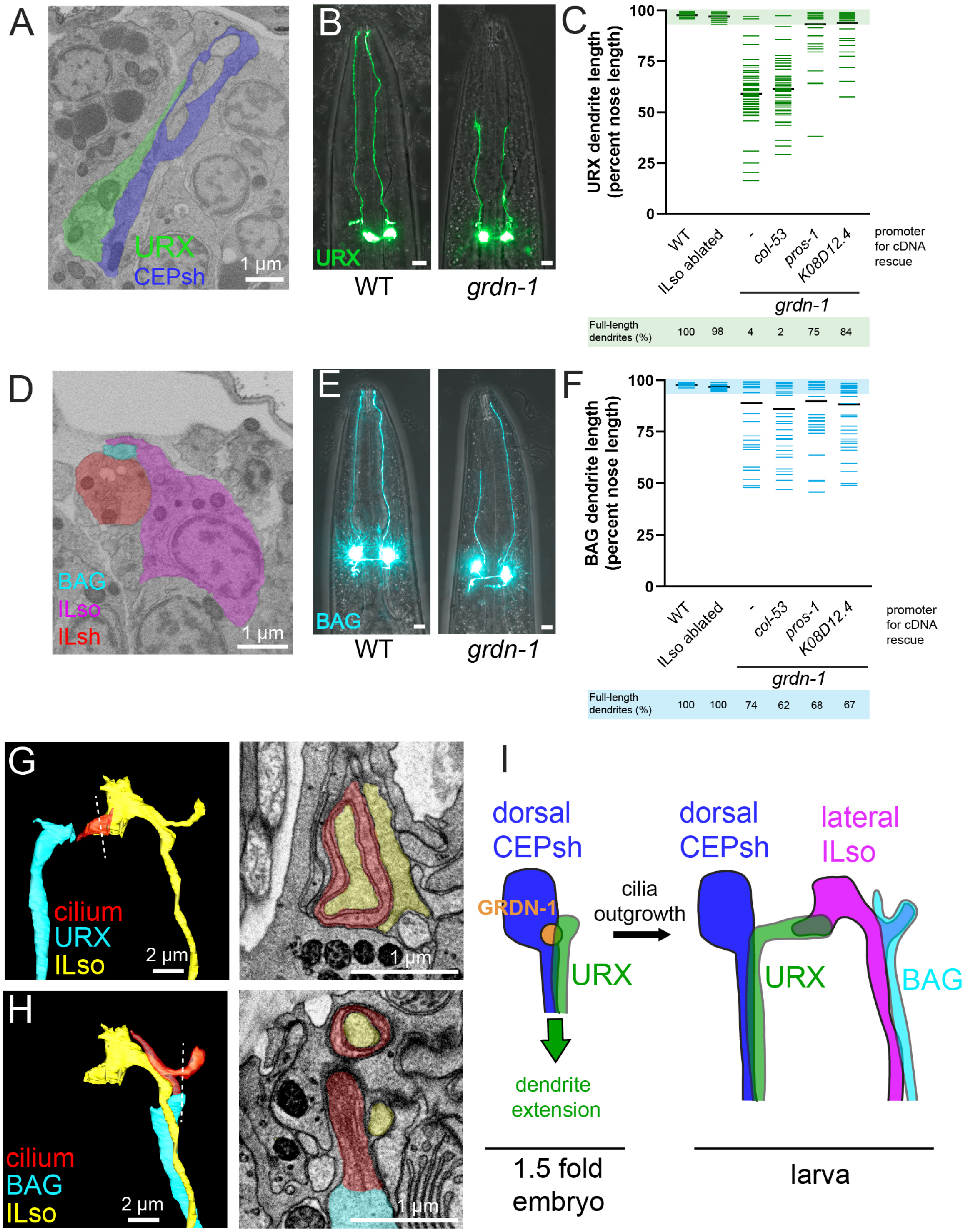


**Supporting Figure S1. URX and BAG dendrites anchor to guidepost glia, not ILso glia**

(A, D) Single EM sections from a comma-stage embryo showing (A) URX (green) and dorsal CEPsh (dark blue) or (D) BAG (light blue), lateral ILso (magenta), and lateral ILsh (red), showing close apposition of URX dendrite ending to dorsal CEPsh and BAG dendrite ending to lateral ILsh at the onset of dendrite extension. (B, E) (B) URX (*flp-8p*) and (E) BAG (*flp-17p*) neurons in wild-type (WT) and *grdn-1(ns303)* animals, showing dendrite extension defects. (C, F) Quantification of (C) URX or (F) BAG dendrite lengths as a percentage of distance from the cell body to the nose (μm); colored bars represent individual dendrites, n≥50 per genotype; black bars represent mean for each genotype. "Full length" is defined as the wild-type mean ± 5 SD (shaded region). Dendrites remain full length in ILso ablated (*col-53*p:Diptheria Toxin A) animals. Expression of wild-type *grdn-1* cDNA in embryonic ILso glia (*col-53*p) in *grdn-1(ns303)* animals does not rescue URX or BAG dendrite extension, whereas expression in embryonic dorsal CEPsh glia (*pros-1p* or *K08D12.4*p) rescues URX but not BAG dendrite extension. Note that prior work also showed that another dendrite anchoring molecule, MAGI-1, rescues URX but not BAG dendrite extension when expressed in embryonic dorsal CEPsh glia (*pros-1p)* (Cebul et al., Development 2026). (G, H) Reconstruction (left) and single section (right) from EM of wild-type L4 larva showing (G) URX cilium (red), URX (blue), and lateral ILso (yellow) or (H) BAG cilium (red), BAG (blue), and lateral ILso (yellow). Note that the same lateral ILso is shown in both cases. Microtubules are visible within the cilia in the EM sections. (I) Schematic of URX development through GRDN-1-dependent anchoring to the dorsal CEPsh glial guidepost cell during dendrite extension in the embryo (left), followed by cilia outgrowth to form the mature attachment to lateral ILso glia observed in larvae (right). URX, green; dorsal CEPsh, dark blue; GRDN-1-dependent anchor site, orange; BAG, light blue; lateral ILso, magenta.

**SUPPORTING FIGURE 2**

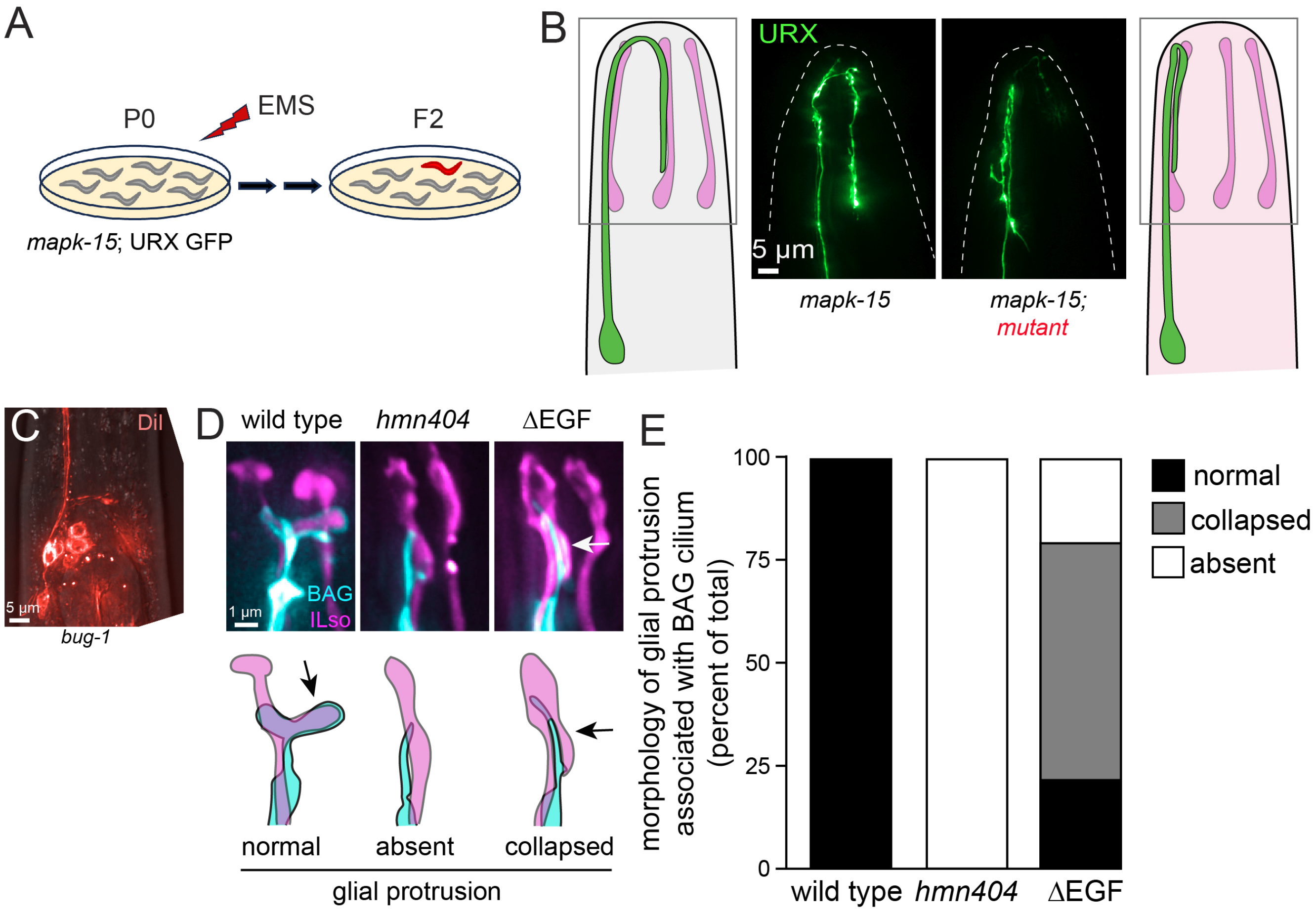


**Supporting Figure 2. BUG-1 is required for cilia-glia attachment**

(A) Schematic of forward genetic screen. The URX overgrowth mutant *mapk-15* was used to increase the length of the URX dendrite ending (McLachlan et al., PLoS Genet. 2018), making it more apparent if the dendrite ending projects to the lateral sensory fascicle (wild type) or remains in the dorsal sensory fascicle (mutant). P0 animals bearing the URX overgrowth mutation *mapk-15(hmn5)* and the URX GFP marker *flp-8*p:GFP were mutagenized with ethyl methanesulfonate (EMS), grown for two generations to produce homozygous mutations (F2), and visually screened in non-clonal pools using a fluorescence stereomicroscope. (B) Diagram and fluorescence micrographs showing typical phenotypes observed in most adult F2s (left) or in rare adult F2s exhibiting the desired mutant phenotype (right). In most adults, the *mapk-15* defect causes the URX dendrite ending to continue to grow along the lateral ILso (left); in animals in which URX fails to project laterally, it makes a sharp Uturn and grows back along the dorsal ILso (right). (C) DiI staining in *bug-1(hmn404)* larva which indicates that cilia are generally intact. (D) Fluorescence micrographs and schematics of BAG dendrite endings (blue) with lateral ILso endings (magenta) in wild type, *bug-1(hmn404)*, and *bug-1(hmn423)* animals (image for wild type is the same as in Figure 2), showing the normal glial protrusion (wild type), absence of a glial protrusion (*bug-1(hmn404)),* or the collapsed glial protrusion pointing posteriorly along the BAG cilium (*bug-1(hmn423),* BUG-1b∆EGF). Arrows indicate glial protrusion. (E) Quantification of ILso glial protrusion morphology in wild-type, *bug-1(hmn404)*, and BUG-1b∆EGF animals; $n \geq 50$ per genotype.

**SUPPORTING FIGURE S3**

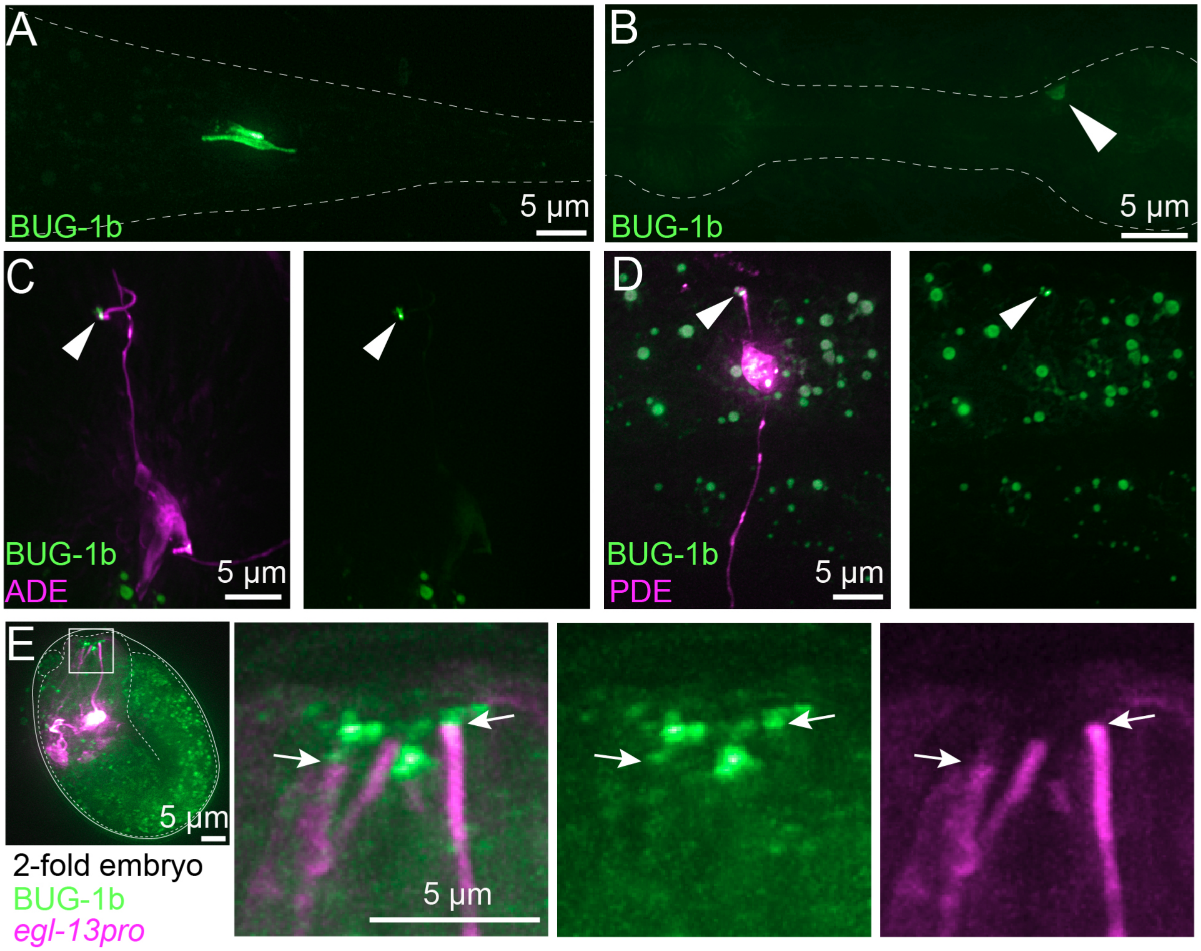


**Supporting Figure S3. BUG-1b localizes to neuron-glia attachments**

(A) BUG-1b-sfGFP localization in the tail, likely corresponding to phasmid neurons PHA and PHB interacting with PHsh glia and the PQR cilium attaching to PHso2 glia. (B) BUG-1b-sfGFP localization in a neuron near the posterior pharyngeal bulb, likely corresponding to the oxygen-sensing neuron AQR, the only ciliated neuron that is not known to interact with a glial partner. Localization may be mostly intracellular in the soma. (C, D) BUG-1b-sfGFP (green; white arrowheads) localizes near the cilium of (C) ADE (magenta, *dat-1p*) and (D) PDE (magenta, *dat-1p*). Other green signal is due to autofluorescent gut granules. (E) BUG-1b-sfGFP in a 2-fold embryo, the earliest stage at which expression was detected. URX and BAG dendrites are shown (arrows; magenta, *egl-13p*). At this stage, cilia have not yet formed. Due to localization of BUG-1b in nearby sense organs, it is ambiguous whether a cap of BUG-1b is present at the URX and BAG dendrite endings at this stage.

**SUPPORTING FIGURE S4**

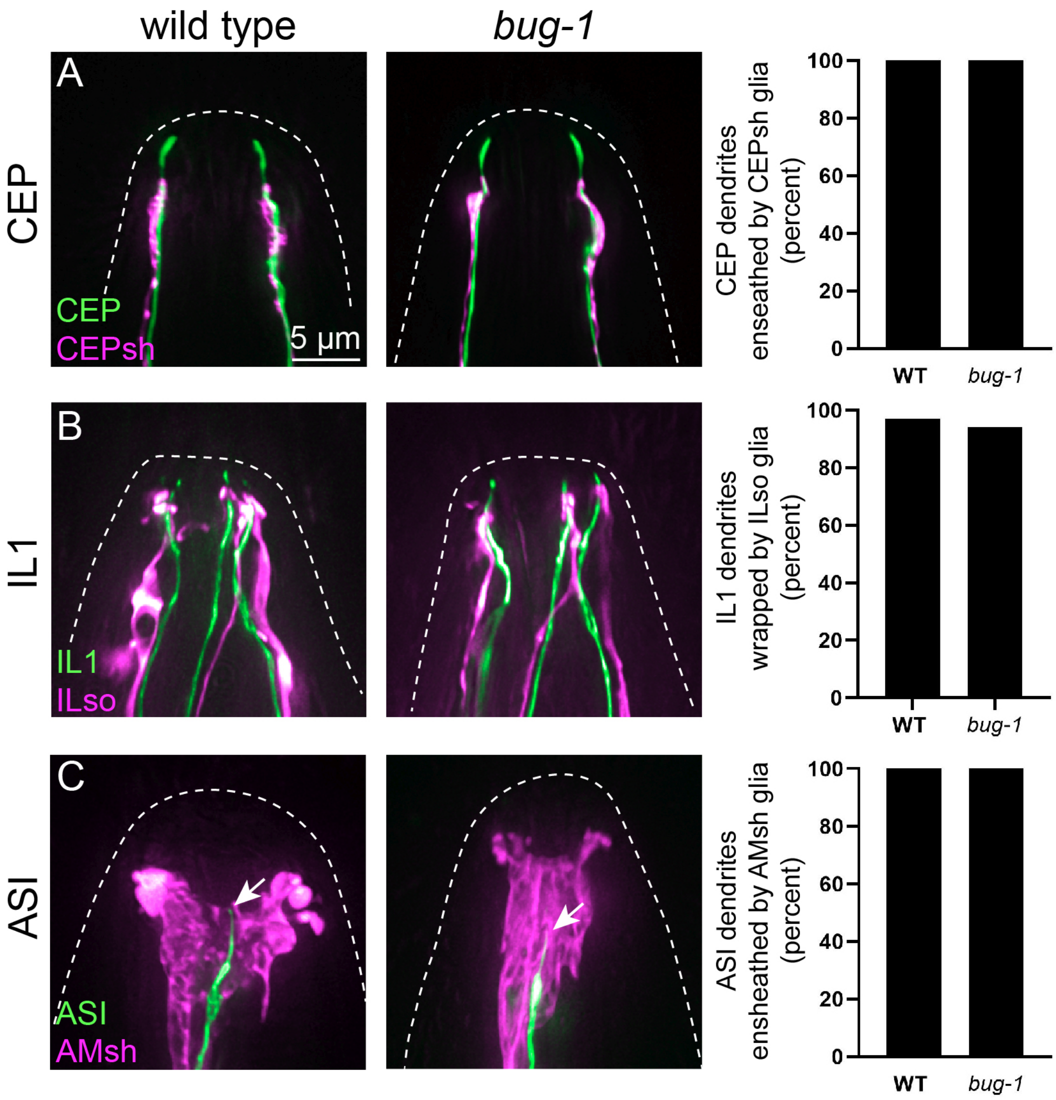


**Supporting Figure S4. Other neuron-glia attachments do not require *bug-1***

(A-C) Images of the dendrite endings and their associated glia in wild-type (left) and *bug-1(hmn404)* (right) animals, with quantification of the fraction of dendrites ensheathed by glia, for (A) CEP (green, *dat-1*p) and CEPsh glia (magenta, *hlh-17*p); (B) IL1 (green, *flp-3*p) and ILso glia (magenta, *grl-18*p); and (C) ASI (green, *daf-7*p) and AMsh glia (magenta, *F16F9.3*p). ASI cilia (arrows) are mispositioned posteriorly relative to the AMsh glial cell in *bug-1*.

**SUPPORTING FIGURE S5**

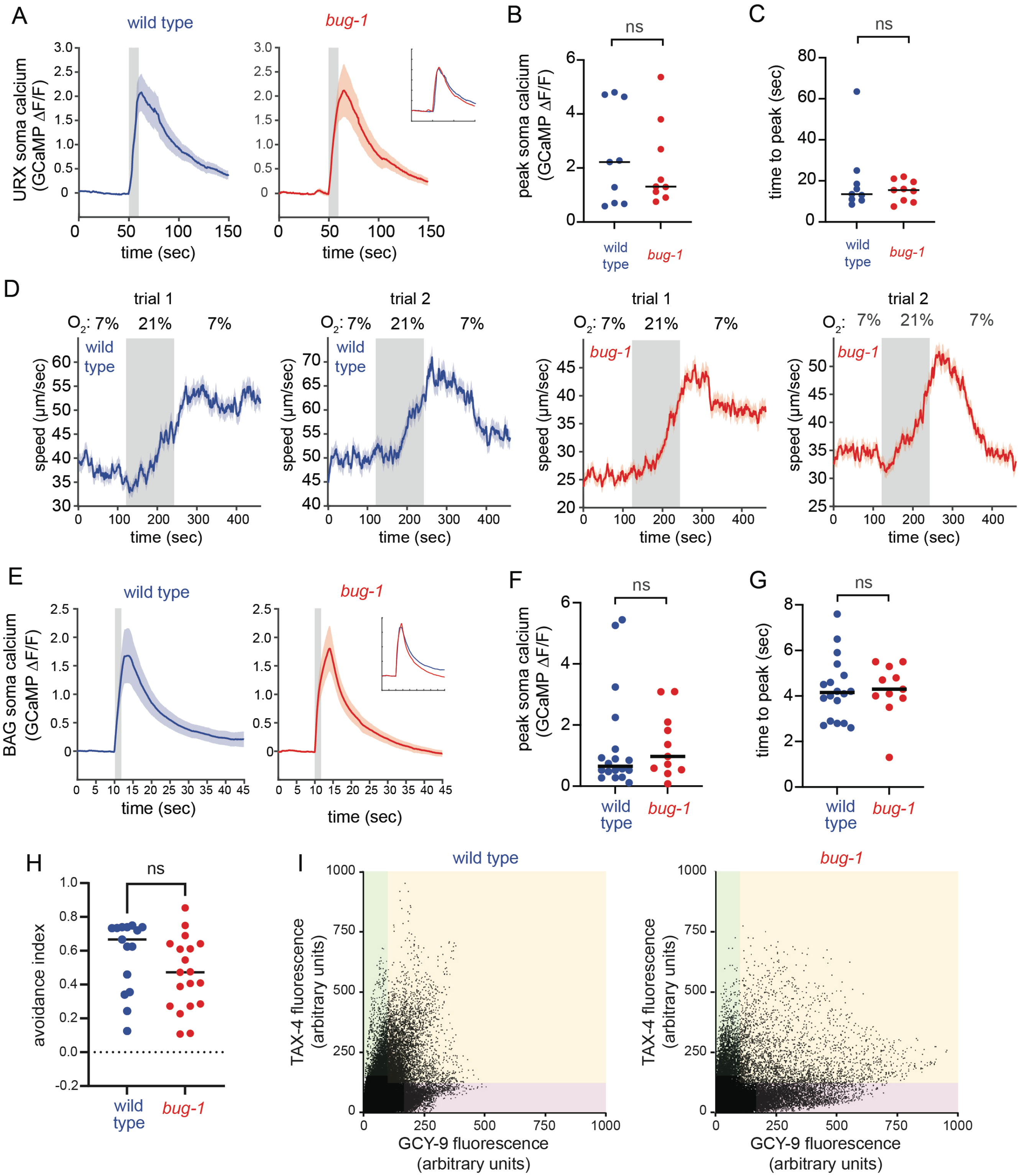


**Supporting Figure S5. The effects of glial attachments on cilia function and organization**

(A) Quantification of change in GCaMP signal in URX cilium. 7% $O_2$ is replaced with ambient (21%) $O_2$ from t=50 sec to t=60 sec (gray shading). Inset shows overlay of wild-type and *bug-*

*1* mean responses. Solid lines, mean; shaded red or blue, SEM. (B) Peak calcium response and (C) time to peak in sec for experiment shown in (A). *P* values from comparisons of wild-type and *bug-1* responses were computed using Student's T test with Welch's correction. *n* = 9 per genotype. (D) Quantification of behavioral responses to $O_2$, measured as change in instantaneous speed in response to shifts from 7% $O_2$ to 21% $O_2$ (gray shading) and return to 7% $O_2$. Populations were adapted to 7% $O_2$ and then subjected to two sequential trials. Both wild-type and *bug-1* mutant animals exhibited increased speed following the first trial, which became the baseline speed in the second trial. Note that the y-axis scales are adjusted to emphasize the overall shapes of the responses, but speeds are consistently higher in the second than first trial for both genotypes and overall lower in *bug-1* than wild type in both trials. Solid lines, mean; shaded red or blue, SEM. (E) Quantification of change in GCaMP signal in BAG soma (compare with change in BAG cilium, Fig. 5C). Experimental conditions are described in Fig. 5C. Inset shows overlay of wild-type and *bug-1* mean responses. Solid lines, mean; shaded red or blue, SEM. (F) Peak calcium response and (G) time to peak in sec for experiment shown in (E). *P* values from comparisons of wild-type and *bug-1* responses were computed using Student's T test with Welch's correction. *n* = 18 wild type, *n* = 11 *bug-1*. (H) Quantification of behavioral responses to $CO_2$, measured as an avoidance index that reflects the fraction of animals on each side of an arena with 10% $CO_2$ on one side and ambient $CO_2$ on the other side. (I) Quantification of relative localization of GCY-9 and TAX-4 in BAG cilia. Scatter plots show fluorescence intensity per voxel of endogenous GCY-9-mApple and TAX-4-GFP11 co-expressed with BAG-specific GFP1-10. Each dot is a voxel from 3D optical stacks of 15 cilia per genotype. Green, red, and yellow regions are drawn arbitrarily to highlight regions containing pixels with high TAX-4 but low GCY-9 (green); low TAX-4 but high GCY-9 (purple); or coexpression of TAX-4 with GCY-9 (yellow).
